# F-actin patches nucleated on chromosomes coordinate capture by microtubules in oocyte meiosis

**DOI:** 10.1101/265009

**Authors:** Mariia Burdyniuk, Andrea Callegari, Masashi Mori, François Nédélec, Péter Lénárt

## Abstract

Capture of each and every chromosome by spindle microtubules is essential to prevent chromosome loss and aneuploidy. In somatic cells, astral microtubules search and capture chromosomes forming lateral attachments to kinetochores. However, this mechanism alone is insufficient in large oocytes. We have previously shown that a contractile F-actin network is additionally required to collect chromosomes scattered in the 70-μm starfish oocyte nucleus. How this F-actin-driven mechanism is coordinated with microtubule capture remained unknown. Here, we show that after nuclear envelope breakdown Arp2/3-nucleated F-actin patches form around chromosomes in a Ran-GTP-dependent manner, and we propose that these structures sterically block kinetochore-microtubule attachments. Once F-actin-driven chromosome transport is complete, coordinated disassembly of these F-actin patches allows synchronous capture by microtubules. Our observations indicate that this coordination is necessary, as early capture of chromosomes by microtubules would interfere with F-actin-driven transport leading to chromosome loss and formation of aneuploid eggs.

## Introduction

Capture of chromosomes by spindle microtubules is an early step of cell division essential for subsequent alignment and segregation of chromosomes by the spindle apparatus. Failure to capture even a single chromosome will delay mitotic progression and may result in aneuploidy, a major cause of carcinogenesis in somatic cells. Since in oocyte meiosis the spindle assembly checkpoint is weakened or absent, in egg cells failure to capture chromosomes ultimately leads to aneuploidy (Shao et al., 2013; Kolano et al., 2012). Aneuploid eggs develop to unviable or severely impaired embryos that in humans is one of the most common causes of infertility and birth defects (Webster and Schuh, 2017).

Mitchison and Kirschner recognized that ‘dynamic instability’, the rapid growth and shrinkage of centrosome-nucleated microtubules is an effective means for microtubules to explore the cellular space in their search for chromosomes (Kirschner and Mitchison, 1986; Mitchison and Kirschner, 1984). The proposed microtubule ‘search-and-capture’ has since been validated in live cells (Rieder and Alexander, 1990; Hayden et al., 1990) and the molecular details of the initial attachments have also been understood. These so-called lateral attachments form between the kinetochore and the microtubule lattice and involve molecular motors, dynein in particular, which transport captured chromosomes pole-ward (Kapoor et al., 2006; Cai et al., 2009; Li et al., 2007; Yang et al., 2007). Subsequently, these lateral attachments are replaced by end-on attachments to allow bi-orientation of chromosomes on the spindle (Shrestha and Draviam, 2013).

Computer simulations by different laboratories recapitulated key features of ‘search-and-capture,’ confirming that this mechanism works effectively within a range of approx. 10 μm, sufficient to capture chromosomes in a typical rounded somatic cell with diameter of approx. 30 μm (Wollman et al., 2005; Holy and Leibler, 1994). However, simulations also predicted that additional ‘facilitation mechanisms’ are required to match the rapid temporal dynamics observed *in vivo* (Wollman et al., 2005). Several such mechanisms have since been identified including selective microtubule stabilization near chromosomes (Carazo-Salas and Karsenti, 2003; Moss et al., 2009; Kaláb et al., 2006), branched microtubule nucleation on microtubules (Petry and Vale, 2015), chromatin-mediated microtubule nucleation (Heald et al., 1996; Karsenti et al., 1984) and microtubule pivoting (Kalinina et al., 2013). Together, these mechanisms render chromosome ‘search-and-capture’ in somatic cells fast and highly efficient (Heald and Khodjakov, 2015).

Oocytes are much larger than somatic cells, as they store nutrients to support early embryonic development. This includes cytoplasmic as well as nuclear components; hence, oocytes not only have a large cytoplasm, but also a large nucleus, historically referred to as the germinal vesicle (Lénárt and Ellenberg, 2002). During meiosis, oocytes divide extremely asymmetrically to retain these stored components in the fertilizable egg. For this reason, across animal species, the meiotic spindle is small and located very eccentrically, anchored to the cell cortex to produce tiny polar bodies (Crowder et al., 2015).

The specific cellular geometry of oocytes, featuring a large nucleus and a small meiotic spindle challenges chromosome ‘search-and-capture’. Indeed, we have shown that in starfish oocytes the known microtubule-driven mechanisms are insufficient, and a contractile actin filament (F-actin) network is additionally required to transport chromosomes to the assembling microtubule spindle (Lénárt et al., 2005; Mori et al., 2011; Bun et al., 2018). The F-actin network forms at nuclear envelope breakdown (NEBD), filling the entire 70-µm diameter nuclear space. It transports chromosomes in approx. 10 minutes to within 35 µm of the microtubule asters located at a cortical position called the animal pole (AP), where the polar bodies will eventually be extruded. Therefore, chromosome congression in starfish oocytes is a two-step process, whereby F-actin-dependent transport delivers chromosomes for capture by spindle microtubules. However, the dynamics of capture by microtubules in this system have not yet been characterized, and whether and how chromosome capture is coordinated with F-actin-driven transport remained unknown.

Here, we tracked chromosomes in 3D at high spatio-temporal resolution in live oocytes to identify individual chromosome capture events. Our data indicate that capture of chromosomes by microtubules needs to be coordinated with F-actin-driven transport, because early capture events would interfere with F-actin-driven transport by causing a local collapse of the F-actin network. We show that this coordination is achieved by Arp2/3-nucleated F-actin patches surrounding chromosomes. These patches form in a Ran-GTP-dependent manner at NEBD, and sterically prevent microtubule-kinetochore attachments for approx. 5 minutes after NEBD. We integrate these results in a computational model, and find that two-step chromosome congression in starfish oocytes is well explained by the classical ‘search-and-capture’ model when we add the single feature of preventing chromosome capture during the initial F-actin-driven transport.

## Results

### Microtubule capture events can be identified on high-resolution chromosome trajectories

Upon entry into meiosis, following the onset of NEBD, a contractile F-actin network forms in the 70-µm nucleus of starfish oocytes and transports chromosomes to the animal pole (AP) (Mori et al., 2011). Starfish oocytes do contain centrosomes, which form two microtubule asters at the AP (Borrego-Pinto et al., 2016a). We have shown earlier by tubulin immunofluorescence that these microtubules extend to 30-40 µm from the AP (Lénárt et al., 2005). Once the F-actin network transports chromosomes within this ‘capture range’, chromosomes are caught and are eventually incorporated into the first meiotic spindle forming at the AP (Lénárt et al., 2005) (Fig. 1A).

**Figure 1.**
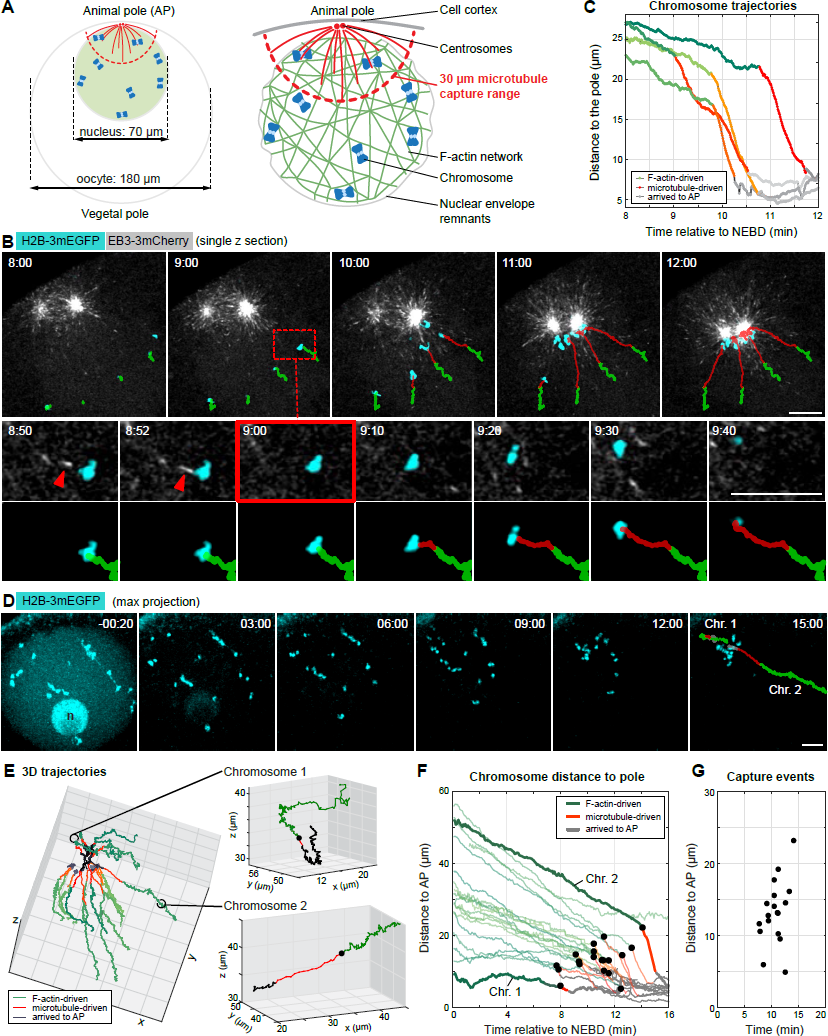
After actin-driven congression, chromosomes form lateral attachments and are transported along microtubules to the spindle poles. **(A)** Left: scheme of an immature starfish oocyte with the nucleus anchored at the animal pole (AP) and with centrosomes nucleating astral microtubules. Right: schematics of the nuclear region after NEBD. The F-actin network fills the nuclear region and as it contracts, transports embedded chromosomes towards the AP. Chromosomes delivered within the capture range of astral microtubules are captured, and transported on microtubules to the centrosomes at the AP. **(B)** Selected frames from a time series of single confocal sections through the nuclear region of an oocyte expressing EB3-3mCherry to visualize microtubule plus-tips (grey) and H2B-3mEGFP to label the chromosomes (cyan). See also Video 1. Chromosome trajectories are overlaid onto the images: green denotes actin-driven transport; microtubule-driven transport is shown in red. Red arrowheads mark the contact between the microtubule and chromosome. Lower panels: zoom of the area marked with a dashed square, selected time points around microtubule capture are shown. **(C)** Plot of distance of chromosomes to the AP over time, calculated from the trajectories shown in **(B). (D)** Selected maximum intensity z-projections from a confocal time series though the oocyte’s nuclear region during chromosome congression. Chromosomes (H2B-3mEGFP) are shown in cyan. n - nucleolus. **(E)** 3D plot of chromosome trajectories derived from the movie shown in **(D)**, with trajectories of one of the furthest and one of the closest chromosomes shown separately. Green: F-actin-driven transport, red: transport on microtubules, black dots: capture events. **(F)** Plot of chromosome distance to the AP over time for the same dataset shown in **(D)** and **(E)**, and labeled as on **(E). (G)** Plot of capture events identified on **(E)**. Scale bars: 10 µm. Time is given as mm:ss relative to NEBD.

In order to visualize individual chromosome capture events in live oocytes, we imaged chromosomes and growing microtubule tips by acquiring single confocal sections at high spatial and temporal resolution, starting from the NEBD until chromosomes are collected at the AP (Fig. 1B, Video 1). We then automatically tracked chromosome motion (Mori et al., 2011; Monnier et al., 2012). By comparison of chromosome trajectories to microtubule dynamics, we could clearly identify individual events of capture: initially the F-actin network transported chromosomes at a lower speed and with the motion being less directed and more diffusive (Fig. 1B, C, green trajectories). Then, shortly (10-30 s) following a visible direct contact to a microtubule, the chromosomes switched to a faster and directed motion (Fig. 1B, C, red trajectories). The speed of motion (9.22±2.86 µm/min) after the switch matches well the speed expected for dynein-driven transport (Barisic et al., 2014). Consistently, microtubule-driven transport is abolished by the dynein small-molecule inhibitor, Ciliobrevin D (Fig. S1A, B). Together, this suggests that chromosomes are caught by microtubules forming canonical lateral kinetochore-microtubule attachments, immediately followed by dynein-driven transport (Cai et al., 2009; Li et al., 2007; Yang et al., 2007).

These observations establish that chromosome capture by microtubules manifests as a switch-like change in the chromosome’s trajectory – a transition from a slow, more diffusive to a fast, directed motion that also often coincides with a change of the overall direction. To systematically identify such capture events, we tracked chromosomes in the entire nuclear volume in 3D, which we were able to achieve at 3 s time resolution (Fig. 1D, E). In these recordings, a chromosome capture event was identified as a time point followed by at least four subsequent steps of unidirectional and fast motion (Fig. 1D, Chromosome 2), or at least two such steps if it coincided with a change in the overall direction (Fig. 1D, Chromosome 1). Every capture event was confirmed by examining the 3D trajectories (Fig. 1E), as well as plots of the chromosome-AP distance over time (Fig. 1F). These latter plots visualize the pole-ward/radial velocity component corresponding to the expected direction of transport on astral microtubules (Fig. 1F). By these stringent criteria, we were able to identify capture events in approx. 50% of chromosome trajectories in an unbiased manner (Fig. S1C). Taken together, by analyzing chromosome trajectories we were able to reliably identify chromosome capture events in the entire 3D nuclear volume of starfish oocytes.

### Chromosome capture is coordinated by an F-actin-dependent mechanism

Being able to identify capture events allowed us to ask how F-actin-driven transport and microtubule capture are coordinated during chromosome congression. To this end, we compared chromosome capture in untreated control oocytes and oocytes, in which F-actin-driven transport was abolished.

To inhibit F-actin-driven transport, we developed a protocol to acutely depolymerize F-actin after NEBD, because, as we had shown earlier, NEBD in starfish oocytes requires F-actin (Mori et al., 2014) (Fig. 2A). Therefore, Latrunculin B (or equal amount of DMSO for controls) was added to the oocytes immediately after NEBD, allowing the formation of the so-called F-actin shell required for the rupture of nuclear membranes (Fig. 2B, 00:30). Thereafter, Latrunculin B rapidly (in 2-3 min) disrupted all F-actin structures and abolished F-actin-driven chromosome transport, resulting in massive loss of chromosomes distal to the AP (Fig. 2B, Video 2). The affected F-actin structures included the cell cortex, the F-actin network in the nuclear region, as well as dense patches of F-actin, which were previously observed to surround chromosomes (Fig. 2B) (Lénárt et al., 2005).

**Figure 2.**
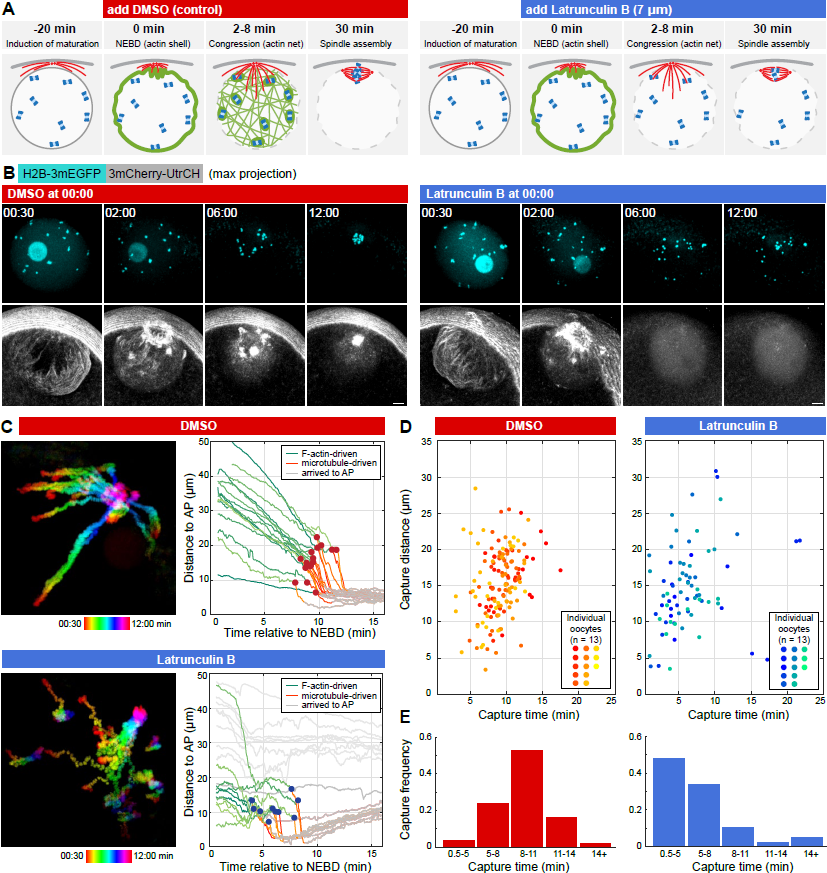
An F-actin-dependent mechanism delays chromosome capture. **(A)** Schematics of the experimental protocol for acute depolymerization of F-actin. **(B)** Selected maximum intensity z-projections from a 3D confocal time series though the oocyte’s nuclear region during chromosome congression. Chromosomes (H2B-3mEGFP) are in cyan and F-actin (3mCherry- UtrCH) in grey. Scale bars: 10 µm. See also Video 2. (**C**) Left: pseudo-colored time projection of a 3D confocal time series of chromosome congression (labeled with H2B-3mEGFP) in control or Latrunculin B treated oocytes. See also Video 3. Right: plot of chromosome distance to the AP over time, for the control and Latrunculin B treated oocytes shown on the left. Labels: green: F-actin-driven transport, red: microtubule-driven transport, grey: arrived to the spindle, dots: chromosome capture events. **(D)** Chromosome capture events identified for 13 pairs of control and Latrunculin B treated oocytes (plotted in a different color for each oocyte). **(E)** Histograms of the data shown in (**D**). Time is given relative to NEBD for all panels.

We next combined this protocol with high resolution tracking of chromosome motion. In control oocytes, trajectories showed the two clearly distinguishable phases of F-actin- and microtubule-driven transport (Fig. 2C, Video 3). The transition between the two phases, i.e. the capture events, occurred rather synchronously (9.21±2.36 minutes after NEBD) showing a slight distance-dependence (Fig. 2C). Upon acute F-actin depolymerization distal chromosomes outside of the microtubule capture range were lost, as expected (Fig. 2B, C, and Mori et al., 2011). However, to our surprise, chromosomes initially positioned within the microtubule capture range were efficiently captured by microtubules, and capture of these chromosomes occurred earlier than in controls (6.16±3.92 minutes after NEBD) (Fig. 2B-D). This difference is clearly visualized on histograms of capture events: the majority of captures occur between 8-11 min after NEBD in control oocytes, in contrast to the peak between 0.5-5 min for Latrunculin B treated oocytes (Fig. 2E).

Thus, in control oocytes capture of chromosomes is prevented in the approx. first 5 minutes after NEBD, whereas when F-actin is depolymerized capture starts immediately after NEBD (Fig. 2D). This implies that besides chromosome transport, F-actin structures play an additional role in coordinating chromosome capture by preventing microtubule capture until F-actin-driven transport is completed.

### The F-actin network is disrupted by chromosomes transported along microtubules

Next, in order to explore the potential function and the underlying mechanism of the F-actin-dependent delay of chromosome capture, we characterized the two prominent F-actin structures present during this relevant time window: the F-actin network and the F-actin patches. Our hypothesis was that capture by microtubules may interfere in some way with transport of chromosomes by the F-actin network, and therefore capture by microtubules needs to be prevented until F-actin-driven transport is complete. We envisaged two scenarios: (i) coordination may be achieved by the F-actin network hindering access of microtubules to chromosomes. (ii) Alternatively, microtubule attachments could form uninhibited, but the F-actin network might physically entrap chromosomes to prevent their pole-ward transport.

To test these possibilities, we first fixed and co-stained oocytes for F-actin and microtubules and imaged them at high resolution in 3D. In these samples we could clearly visualize astral microtubules penetrating the F-actin network (Fig. 3A). Furthermore, we could not detect inhomogeneities in the microtubule density, suggesting that microtubules can grow just as freely into the F-actin network as in the cytoplasm (Fig. 3A). Additionally, we monitored microtubule dynamics in live oocytes either treated with Latrunculin B or DMSO, using the protocol above (Fig. 2A). Quantification of microtubule lengths showed no significant difference between Latrunculin B and DMSO-treated oocytes (Fig. 3B and S3A). Based on these results we concluded that the F-actin network does not present an obstacle to microtubule growth, and therefore the F-actin network *per se* does not appear to restrict access of microtubules to chromosomes.

**Figure 3.**
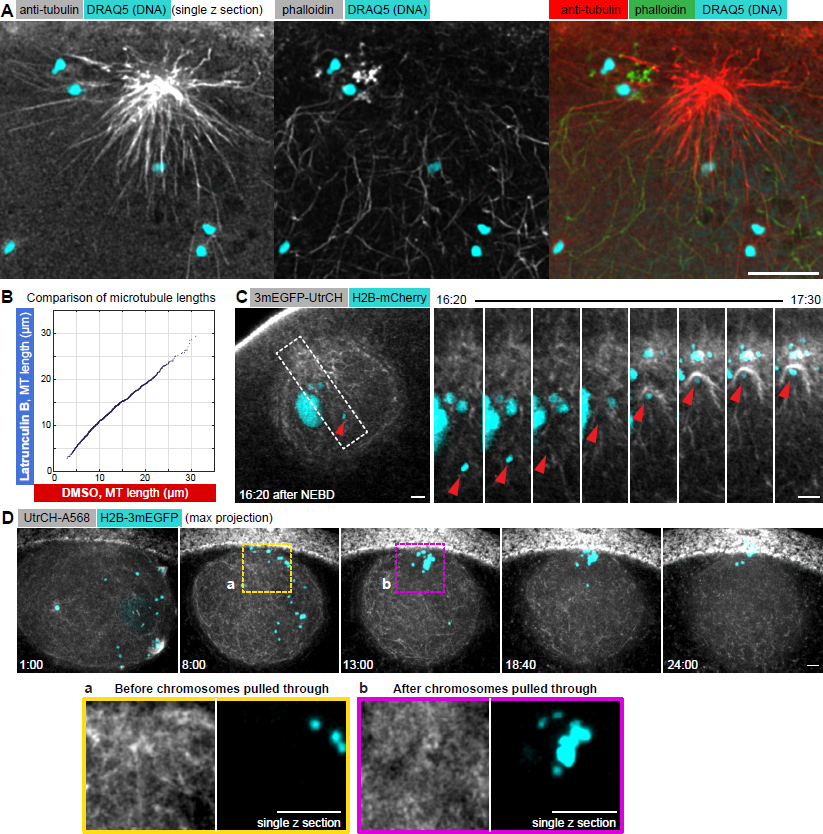
The F-actin network does not prevent chromosome capture and transport by microtubules, but transport along microtubules interferes with F-actin network integrity. **(A)** A single selected slice from a deconvolved confocal stack of an oocyte fixed 5 minutes after NEBD and stained for tubulin (red), F-actin (green) and for chromosomes using Draq5 (cyan). **(B)** Microtubule length distribution measured in control and Latrunculin B treated oocytes and plotted against each other. **(C)** Selected single confocal sections acquired over time showing the nuclear area of an oocyte expressing 3mEGFP-UtrCH (grey) and H2B-mCherry (cyan). See also Video 4. Right: zoom-in on the region marked by a dashed rectangle of a chromosome transported along a microtubule (red arrowhead) causing local collapse of the F-actin network. **(D)** Maximum intensity z-projections of a 3D confocal time series through the nuclear region of an oocyte expressing H2B- mCherry (cyan) and injected with UtrCH-A568 (grey). Single Z-slice zooms of the regions marked by dashed rectangles are shown below visualizing the disruption of the F-actin network where chromosomes are pulled through. Scale bars: 10 µm; 5 µm for the zoom-in image on **(C)**. Time is given as mm:ss relative to NEBD.

We then wondered whether the F-actin network may retain chromosomes by direct binding between chromatin and F-actin filaments, thereby preventing their dynein-driven transport along microtubules. Therefore, we generated chromatin fragments by treating oocytes with Zeocin, which introduces DNA double strand breaks (Fig S3B). Small fragments (approx. 0.5 µm) were sieved through the F-actin network and diffused freely in the nucleoplasm, while large fragments were transported by the network and incorporated into the spindle (Fig. S3B), remarkably similar to the behavior of inert beads of comparable sizes, as shown earlier (Mori et al., 2011). This suggests that chromatin does not directly interact with filaments of the F-actin network.

Next, we tested whether the F-actin network could potentially hinder chromosome transport along microtubules. To this end, we recorded movies with chromosomes and F-actin co-labeled, and we could clearly visualize chromosomes being captured and pulled through the still intact F-actin network, dragging F-actin bundles along (Fig. 3C, Video 4). Further, chromosome capture and transport occurred efficiently even when filaments in the network were strongly stabilized by phalloidin (Fig. S3C). These results indicate that the F-actin network is not sufficient to resist chromosomes transported along microtubules. To the contrary, the movies showed that such pulling through of chromosomes by microtubules causes a local collapse and disruption to the F-actin network, ‘Clearing out’ the network where chromosomes pass (Fig. 3D).

Based on these findings, we conclude that the F-actin network does not present an obstacle for microtubules to grow towards, capture and transport chromosomes pole-ward. Importantly, however, the above observations provide an explanation for why capture by microtubules needs to be coordinated with F-actin-driven transport: if capture started immediately after NEBD, this would lead to local disruption of the F-actin network by early-capture chromosomes being pulled through, interfering with transport of distal chromosomes.

### Disassembly of the F-actin patches coordinates capture by microtubules

Next, we tested the possibility whether the F-actin patches surrounding chromosomes may be responsible for delaying chromosome capture until transport by the F-actin network is complete.

We first characterized the morphology of the F-actin patches and the molecular factors involved in their formation. F-actin patches appear as structures composed of spots of dense F-actin, reminiscent of endocytic sites or other similar Arp2/3 nucleated structures (e.g. Kaksonen et al., 2003), clearly distinct from the F-actin network composed of long filament bundles (Fig 4A). We detected patches of variable size and intensity on chromosomes, with peripheral chromosomes in contact with nuclear envelope membranes being surrounded by much larger and brighter patches, as compared to those located deeper in the nuclear volume (Fig. 4A).

**Figure 4.**
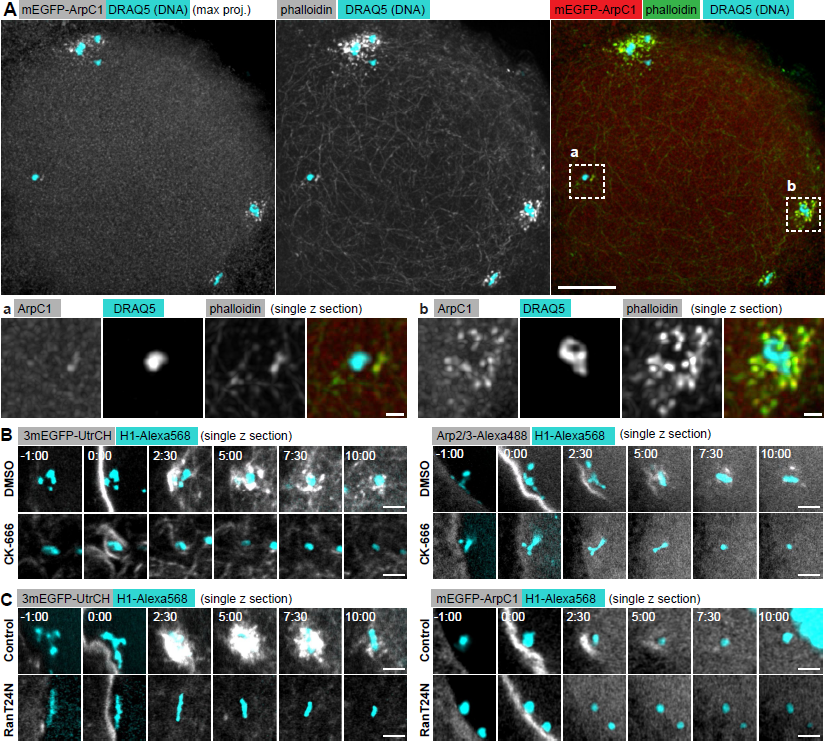
F-actin patches are nucleated on chromosomes by the Arp2/3 complex in a Randependent manner. **(A)** Maximum projection of selected z-sections from a confocal z-stack of an oocyte expressing mEGFP-ArpCl fixed 5 minutes after NEBD and immunostained. anti-GFP was used to enhance mEGFP-ArpC1, Phallodin-A568 to stain F-actin and Draq5 for DNA. Below: selected single z-slices zooming in on F-actin patches marked by dashed rectangles on the overview. Scale bars: 10 µm and 1 µm, respectively. **(B)** Single confocal slices selected from a time series of an oocyte injected with Hl- Alexa568 (cyan) and expressing either 3mEGFP-UtrCH (grey) to label F-actin or injected with Arp2/3- Alexa488 protein (grey) to visualize the Arp2/3 complex. A region around a selected chromosome is shown. Oocytes were treated with CK-666 or with equal amount of DMSO lh before maturation. **(C)** Single confocal slices selected from a time series of an oocyte injected with H1-Alexa568 (cyan) and either 3mEGFP-UtrCH mRNA to visualize F-actin or Arp2/3-Alexa488 protein to visualize the Arp2/3 complex. A region around a selected chromosome is shown. Oocytes were injected with RanT24N protein or equal amount of buffer as control. Time is given as m:ss relative to NEBD. Scale bars: 5 µm.

Consistent with their morphology, F-actin patches were specifically labeled by mEGFP-ArpC1, a subunit of the Arp2/3 nucleator complex (Fig. 4A). Indeed, the small-molecule Arp2/3 inhibitor, CK- 666 blocked recruitment of Arp2/3 and effectively prevented the formation of F-actin patches, while leaving the F-actin network largely intact (Fig. 4B) (Mori et al., 2014). These data thus allow us to conclude that the F-actin patches are nucleated by the Arp2/3 complex, while filaments of the F-actin network are polymerized by other factors, likely formins (Bun et al., 2018). We have shown previously that the F-actin patches are nucleated around DNA-coated beads (Lénárt et al., 2005). This further suggested the involvement of the small GTPase, Ran by analogy to Ran- and Arp2/3-mediated actin nucleation reported in mouse oocytes (Deng et al., 2007). To test this hypothesis, we injected a large amount of RanT24N, a mutated version of Ran locked predominantly in its inactive, GDP-bound form (Dasso et al., 1994). RanT24N injection abolished F-actin patches, indicative of a Ran-GTP- and Arp2/3-dependent nucleation pathway (Fig. 4C).

Next, we quantified the assembly and disassembly kinetics of F-actin patches and correlated this with chromosome capture. To this end, we co-labeled chromosomes and patches (using mEGFP-ArpC1 for Arp2/3 or 3mCherry-UtrCH for F-actin), and imaged these in 3D at high resolution in live oocytes (Fig. 5A, Video 5). We then tracked chromosomes as above and, using chromosome coordinates as reference points, quantified the total patch intensity in a 2.5-µm-radius sphere around every chromosome for each time point (Fig. 5C, G-H). These measurements showed that F-actin patches assemble within 1-2 minutes after NEBD on all chromosomes (with peripheral chromosomes followed by those deeper in the nucleus). The intensity of the patches peaked at around 5 minutes, followed by disassembly, with mEGFP-ArpC1 and 3mCherry-UtrCH intensities dropping to background levels approx. 8 minutes after NEBD (Fig. 5C, G-H).

**Figure 5.**
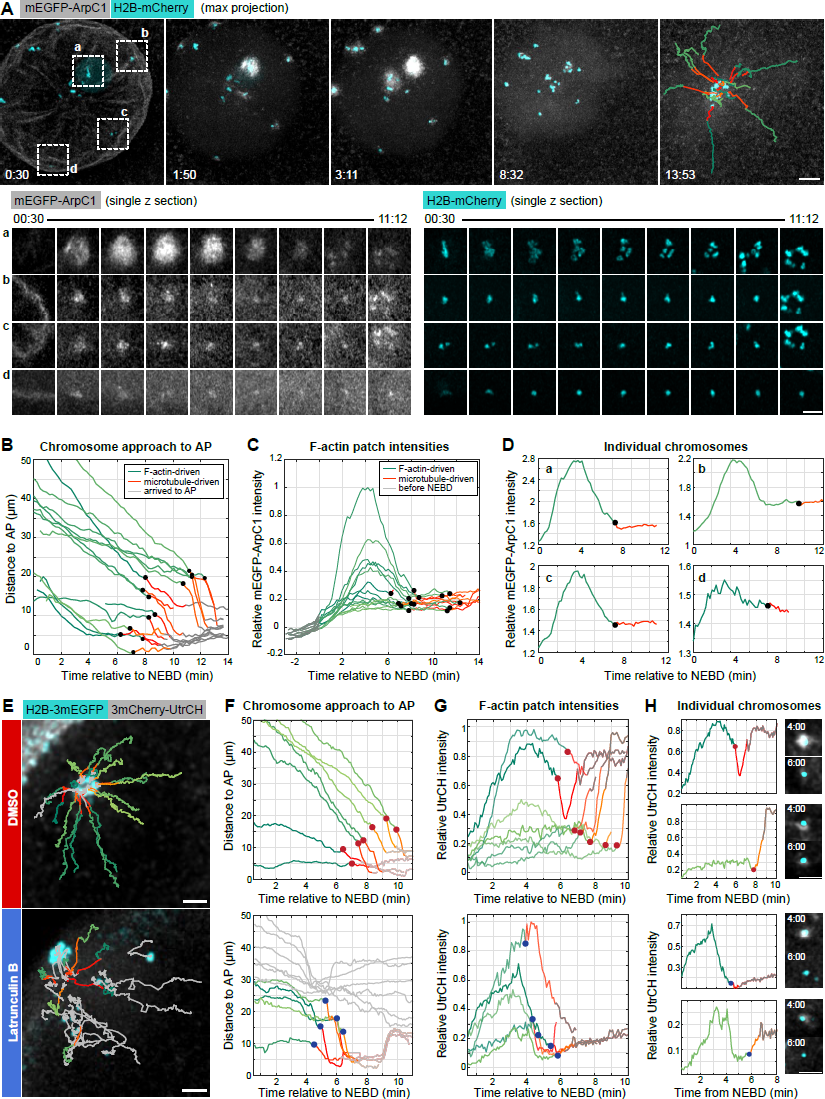
Disassembly kinetics of F-actin patches is tightly correlated with chromosome capture by microtubules. **(A)** Maximum intensity z-projection of selected time points from a time series of the nuclear area of an oocyte expressing mEGFP-ArpCl to label the Arp2/3 complex (grey), and H2B-mCherry to label chromosomes (cyan). Last frame: chromosome tracks overlaid, color-coded as below. See also Video 5. Below: single confocal slices of selected chromosomes marked by dashed rectangles on the overview. Scale bars: 10 µm and 5 µm, respectively. Time is given as mm:ss relative to NEBD. **(B)** Plot of chromosome distance to the AP over time, calculated from the trajectories shown in (**A**). Trajectories are color-coded for actin- (green) and microtubule- (red) driven transport phases and arrival at the spindle (grey). Chromosome capture events are represented by black dots. **(C)** Normalized mEGFP-ArpCl intensity profile for each chromosome tracked in **(B)**. Intensity is calculated in a 5 µm diameter sphere around the chromosome’s center of mass and normalized to the background level before NEBD onset (in grey). Plots are color-coded as in **(B). (D)** Individual plots for chromosomes shown in **(A)**. **(E)** Maximum z-projection of the last time point from a time series of the nuclear area of oocytes expressing 3mCherry-UtrCH to label F-actin (grey), and H2B-3mEGFP to label chromosomes (cyan). Oocytes were treated with DMSO or Latrunculin B, respectively. Chromosome tracks overlaid, color-coded as above. **(F)** Plot of chromosome distance to the AP over time, calculated from the trajectories shown in **(E). (G)** 3mCherry-UtrCH intensity profiles for each chromosome tracked in **(F)**. **(H)** Individual plots for chromosomes shown on **(G)**. Right: single confocal slices of selected chromosomes plotted. Scale bars: 10 µm.

The disassembly kinetics of F-actin patches was remarkably coordinated and largely independent of initial patch size and intensity (Fig. 5A, D). Strikingly, this kinetics of F-actin patch disassembly matched well the time of chromosome capture by microtubules (Fig. 5B, F): the first capture events were detected 5-8 minutes after NEBD, when the first patches disassemble, and most chromosomes were captured very soon after F-actin patch disassembly, 8-11 minutes after NEBD. Notably, at later times a second F-actin accumulation was detected in the spindle area, possibly related to the F-actin spindle observed recently in mouse oocytes (Mogessie and Schuh, 2017). Therefore, for some chromosomes the UtrCH intensity peaks a second time when chromosomes congress at the AP (Fig. 5G, H).

Further substantiating our hypothesis, the correlation between F-actin patch disassembly and chromosome capture also persisted in oocytes in which F-actin patches were acutely depolymerized. After Latrunculin B addition, we observed a complete disassembly of F-actin patches at approx. 4 min after NEBD (slightly delayed compared to the F-actin network) (Fig. 5E, F, Fig. S5). In these oocytes, chromosome capture events closely followed the decline in patch intensities (Fig. 5E-H).

Taken together, patches are F-actin structures distinct from the network and nucleated by the Arp2/3 complex on chromosomes in a Ran-GTP-dependent manner. Quantitative analysis of their disassembly kinetics strongly suggests that F-actin patches are responsible for preventing chromosome capture in the first approx. 5 minutes after NEBD by sterically blocking microtubule-kinetochore attachments.

### ‘Search-and-capture’ extended by an early block recapitulates chromosome congression dynamics

Finally, in order to test potential mechanisms of the F-actin dependent delay in chromosome capture, we integrated our observations in a computational model using the Cytosim software (Nedelec and Foethke, 2007). Within the realistic 3D geometry of the starfish oocyte, microtubules were nucleated from centrosomes located at the AP with dynamics based on dynamic instability and parameters estimated experimentally (Fig. 6A, S6A-E, Video 6) (Magidson et al., 2015; Mitchison and Kirschner, 1984; Wollman et al., 2005).

**Figure 6.**
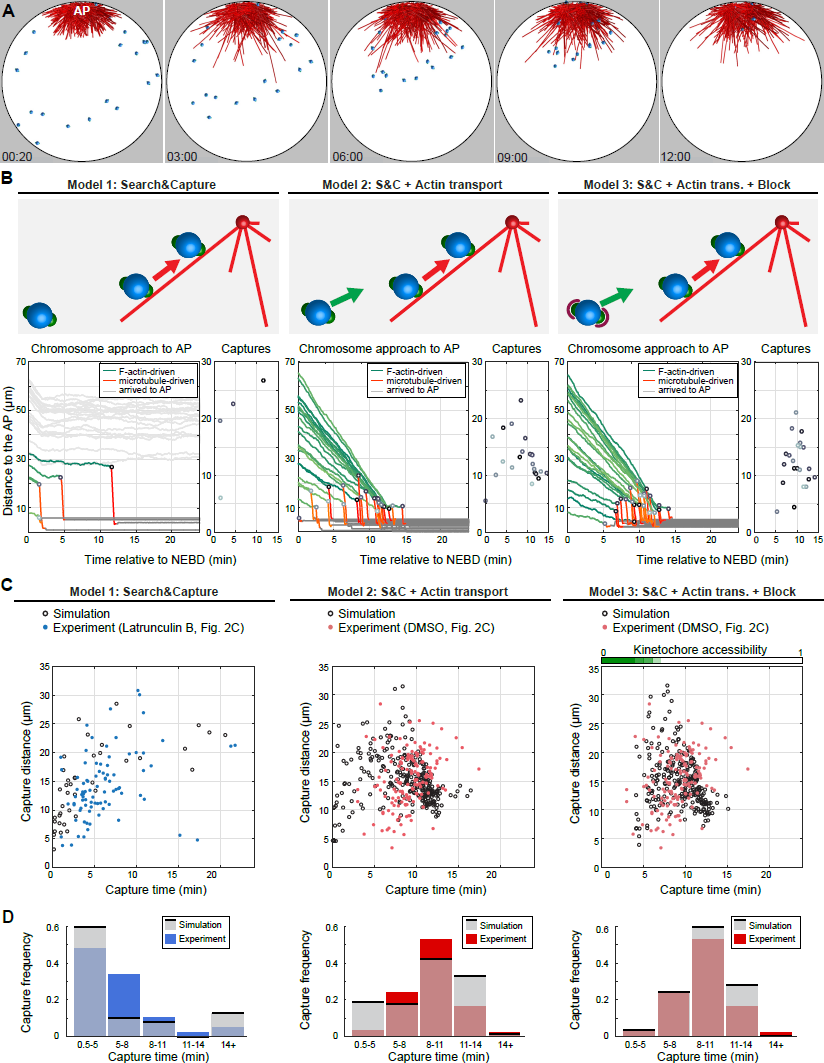
‘Search-and-capture’ expanded by F-actin-driven transport and block of early capture explains the chromosome capture dynamics. **(A)** Renderings of 3D Cytosim simulations of two-staged chromosome congression. Microtubules are in red, chromosomes are blue spheres with green kinetochores. Time starts at NEBD. See also Video 6. **(B)** Schematics of the different models and corresponding plots of chromosome distance to the AP over time and capture events. Trajectories are color-coded for actin- (green) and microtubule- (red) driven transport phases and arrival at the spindle (grey). Chromosome capture events are represented by grey dots. **(C)** Comparison of simulated and experimental chromosome capture dynamics (observed in oocytes, treated with Latrunculin B or DMSO, see Fig. 2). Data from l3 oocytes are shown for both simulations and experiments. **(D)** Plots of individual experimental and simulated capture events and histograms of the same data.

We first simulated the classic ‘search-and-capture’, without F-actin driven transport. This model very closely recapitulated the capture kinetics observed in Latrunculin B treated oocytes: distal chromosomes outside of the capture range (approx. 30 µm) were lost, while chromosomes within the capture range were captured one after the other by random search (Fig 6B-D, Model 1). For chromosomes located close to the centrosomes, capture started immediately after NEBD. In the second model, we included a force field simulating the F-actin network that transports chromosomes towards the AP (Fig. 6B-D Model 2, Fig, S6C). In this scenario, all 22 chromosomes were successfully captured by microtubules within approx. 15 min, with similar kinetics to experiments. However, unlike in experiments, the first capture events occurred immediately after NEBD. Therefore, in the third model we introduced an additional feature to simulate F-actin patches. We set the binding rate of kinetochores to microtubules for the first 4 minutes after NEBD to 0 (no binding) and gradually increased it to 10-30-70-100%, every minute, 4-8 minutes after NEBD, matching the disassembly kinetics of patches. Simulations with this feature added faithfully recapitulated chromosome capture dynamics observed experimentally (Fig. 6B-D Model 3).

Taken together, simulations show that the classical ‘search-and-capture’ model can in principle explain chromosome capture in starfish oocytes. There are only two additional, F-actin-dependent features to be added: (i) transport of distal chromosomes by the F-actin network; (ii) F-actin patches that delay capture of chromosomes until the transport by the F-actin network is complete, and thereby ensure that early capture events do not interfere with transport by the F-actin network.

## Discussion

Taken together, our results evidence a novel mechanism to coordinate chromosome capture at the early steps of spindle assembly. This mechanism is independent of microtubules, and is mediated by F-actin patches nucleated on chromatin by the Arp2/3 complex in a Ran-GTP-dependent manner. We propose that these F-actin patches sterically block microtubule-kinetochore attachments until their synchronous disassembly, thereby coordinating chromosome capture. We show that in starfish oocytes, in which chromosomes are first transported by an F-actin network and then by microtubules, this coordination is necessary to ensure that F-actin driven transport is complete before capture by microtubules begins. Our observations suggest that without this coordination early-captured chromosomes would locally collapse of the F-actin meshwork, and thereby interfere with F-actin-driven transport of distally located chromosomes, leading to chromosome loss.

There are number of common features of oocyte meiosis shared across animal species strongly suggesting that such coordination of chromosome capture is generally required to support these divisions. Animal oocytes store nutrients for the early embryonic development, hence the oocyte as well as its nucleus are exceptionally large (Lénárt and Ellenberg, 2002). In order to retain these stored nutrients for the fertilizable egg, meiotic divisions are extremely asymmetric, and thus the meiotic spindle is small and located near the cell cortex across animal species (Crowder et al., 2015). This specific size and organization necessitates additional mechanisms to collect chromosomes scattered in the large nuclear volume, and thus mechanisms to coordinate capture with these processes. Furthermore, in many species meiotic divisions are coupled to fertilization, and therefore these divisions have additional arrest points that again may require mechanisms to coordinate chromosome capture with other cellular events. Importantly, as the spindle assembly checkpoint is weak or inactive in oocytes, chromosome capture and its coordination have a particular importance in meiosis, as this process is required to produce euploid eggs, and thereby essential for sexual reproduction of the species. Intriguingly, actin structures around chromosomes have been reported in early steps of oocyte meiosis in species ranging from jellyfish (Amiel and Houliston, 2009), through tunicates (Prodon et al., 2006), to *Xenopus* (Yamagishi and Abe, 2017) and mouse (Mogessie and Schuh, 2017). It will be exciting to see in the future whether actin has a similar, essential functions in coordinating chromosome capture in these species as well.

Thus, mechanisms coordinating chromosome capture are very likely required during the specialized oocyte divisions to prevent formation of aneuploid eggs. However, similar mechanisms may play a role in mitosis of somatic cells as well. Although completing ‘search and capture’ as quickly as possible may intuitively appear the optimal solution, delaying chromosome capture and coordinating it with other cellular events is likely to provide advantages, and may contribute to preventing chromosome loss and aneuploidy also in mitosis. Remarkably, in somatic cells chromosomes are pre-arranged in a specific spatial configuration in prometaphase, before capture by microtubules, and this has been shown to accelerate the spindle assembly (Magidson et al., 2011). This suggests that mechanisms may exist also in somatic cells to delay capture of chromosomes until they are favorably positioned for rapid and efficient spindle assembly.

In terms of the underlying molecular mechanism, data from mouse oocytes suggest that Ran-GTP-mediated activation of the Arp2/3 complex is a mechanism conserved across species: in mouse oocytes, chromatin induces the formation of the so called ‘actin cap’, an Arp2/3-nucleated thickening of the cell cortex in a Ran-GTP-dependent manner (Deng et al., 2007). Ran-GTP has been proposed to act through Cdc42 and N-WASP in this case (Dehapiot et al., 2013). If this pathway was broadly present in animal cells, it would explain nucleation of actin on chromosomes when cytoplasmic actin monomers enter the nuclear area upon NEBD – in direct analogy to Ran-mediated activation of spindle assembly factors (Hetzer et al., 2002). Therefore, it will be very interesting in the future to explore the molecular details and conservation of the Ran-GTP pathway leading to Arp2/3 activation, how it is controlled in space and time, and how it may contribute to coordinating chromosome capture in other systems.

Finally, an important remaining question is how such dense F-actin structures can prevent chromosome capture by microtubules. One possibility is a biochemical regulation mediated by molecules that interact with both cytoskeletal systems, many of which have been identified recently (e.g. Chesarone et al., 2010; Henty-Ridilla et al., 2016). However, we are not aware of a specific molecular candidate for F-actin-dependent microtubule disassembly. Therefore, we would favor an alternative and not mutually exclusive mechanism, whereby the dense, branched F-actin network nucleated by Arp2/3 constitutes a physical barrier to microtubule growth (unlike formin nucleated bundles that are expected to bend away and organize networks occupying a relatively smaller volume fraction). Indeed, it is observed in different contexts and cell types, particularly striking in axon growth cones, that the Arp2/3-nucleated F-actin networks strictly exclude microtubules, resulting in sharp separation between the two zones (Lowery and Van Vactor, 2009). It will be very exciting to test this hypothesis in the future, and to explore the diverse physiological functions exclusion of microtubules by branched F-actin networks may have.

## Materials and methods

### Oocyte collection, maturation and injection

Starfish *(Patiria miniata)* were obtained from Southern California (South Coast Bio-Marine LLC, Monterey Abalone Company or Marinus Scientific Inc). Animals were maintained in seawater aquariums at 16°C at EMBL’s Marine Facility. Oocytes were isolated, and mRNAs and other fluorescent markers were injected into the oocytes using mercury-filled microneedles, as described previously (Jaffe and Terasaki, 2004; Borrego-Pinto et al., 2016b). mRNA was injected 24-48 hours before to allow protein expression, while fluorescently labeled protein markers were injected a few hours prior imaging. Oocytes were induced to enter meiosis by addition of 1-methyladenine (1-MA, 10 µM, Acros Organics). NEBD normally initiated 25 minutes after 1-MA addition and only those oocytes, which started NEBD within 40 minutes were considered for analysis.

### Live-cell fluorescent markers

H2B-3mEGFP and H2B-mCherry, EB3-3mCherry, mEGFP-ArpC1, 3mEGFP-UtrCH and 3mCherry-UtrCH were subcloned to the pGEM-HE vector (Borrego-Pinto et al., 2016b). mRNA was synthesized *in vitro* from the linearized DNA template using the AmpliCap-Max T7 High Yield Message Maker kit (Cellscript), followed by polyA-tail elongation (A-Plus Poly(A) Polymerase Tailing Kit, Cellscript). mRNAs were dissolved in water (typical concentration 3-5 µg/µl) and injected into the oocytes up to 5% of the oocyte volume. Histone H1 and Utrophin CH domain (UtrCH) (Burkel et al., 2007) were labeled with Alexa fluorophores as described previously (Bun et al., 2018). Phalloidin labeled with the indicated Alexa fluorophores (Invitrogen) was dissolved in methanol, and was then air-dried prior use and dissolved in PBS for microinjection and immunostaining. For phalloidin, microinjection was performed 2 minutes after NEBD into the nuclear area. H1-Alexa568, UtrCH-Alexa568, Arp2/3- Alexa488, mEGFP-Ndc80 proteins were injected into the oocytes prior to maturation. RanT24N (final concentration approx. 15 μM) protein was injected right before NEBD.

### Drug treatments

For all inhibitor treatments oocytes were first transferred to an Ibidi dish (cat# 80131). In order to acutely depolymerize F-actin, Latrunculin B (final concentration 7 µM) was added directly on the microscope stage in form of a double concentrated solution to the equal volume of seawater contained in the chamber. Nocodazole (final concentration 3.3 µM) and Cytochalasin D (final concentration 40 µM) were diluted from DMSO stocks in seawater and added simultaneously with 1- MA. Ciliobrevin D (final concentration 150 µM) was added 10 minutes prior NEBD. Oocytes were incubated with CK-666 (final concentration 0.5 mM) for 1 h prior maturation. Zeocin (final concentration 100 mg/ml) was added 3.5 h prior maturation. In all cases, control oocytes were treated at the same times with the corresponding amount of the DMSO solvent.

### Immunostaining

Oocytes were fixed at desired times by the fixative composed of 100 mM HEPES pH 7.0, 50 mM EGTA, 10 mM MgSO_4_, 0.5% Triton-X100, 1% formaldehyde, 0.1% glutaraldehyde, as described in (Strickland et al. 2004). Samples were additionally treated with ImageIT (ThermoFisher Scientific) to reduce unspecific antibody binding, and mounted with the antifade agent ProLongGold (ThermoFisher Scientific) between single layers of double-sided adhesive tape (Scotch). Microtubules were visualized by an α-tubulin antibody (DM1α, Sigma-Aldrich, 1:400), and goat-anti-mouse Alexa 488 or Alexa 568 secondary antibodies (1:500). mEGFP-ArpC1 signal was enhanced by an α-GFP antibody (Abcam, ab6556, 1:400). Fixed oocytes were imaged according to Niquist criteria (pixel size: 38 nm, Z-step: 130 nm).

### Image acquisition and processing

Live cell movies were acquired on a Leica SP5 confocal microscope using a 40× HCX PL AP 1.10 NA water immersion objective lens (Leica Microsystems). Fixed samples were imaged on a Leica SP8 microscope equipped with the HC PL APO 1.40 NA 100× oil immersion objective. Where indicated, images were deconvolved using the Huygens software (Scientific Volume Imaging). Fast acquisition rate images were acquired on a Zeiss LSM 880 confocal microscope equipped with the AiryFast module and 40× C-Apochromat LD 1.1 NA water immersion objective lens (typical settings to image a volume of 70×70×60 µm volume: 306×306 pixels (pixel size 224 nm) in *xy* and Z step of 1.4 µm allowing a time resolution of one stack every 3 seconds). AiryFast pixel re-assignment and deconvolution was performed in the Zeiss Zen Black software. Imaging was performed at controlled temperature (19-21°C).

Unless specified, images were loaded and adjusted for brightness and contrast, projected (maximum intensity projection and/or temporal color code for time projections) and filtered (Gaussian blur, typically 0.5-1 pixels) in Fiji/ImageJ (Schindelin et al., 2012). Chromosome tracking was performed in 3D using either a custom Matlab routine (Monnier et al., 2012; Mori et al., 2011), or Imaris (Bitplane). Chromosome capture events were identified manually by combining a number of quantitative measures and by examining the 3D trajectories, as well as plots of the chromosome-AP distance over time. Specifically, a chromosome capture event was identified as time point followed by at least four subsequent steps of unidirectional and fast motion, or at least two such steps if it coincided with a change in the overall direction. To calculate the mEGFP-ArpC1 intensity around chromosomes, we tracked chromosomes in Imaris and used a custom XTension script in Matlab to define a spherical volume around chromosome and measured the intensity contained within. Analysis of tracks was performed and plots were generated in Matlab (MathWorks). All figures were assembled in Adobe Illustrator CS6.

### Simulations

Chromosome congression was modeled in the Cytosim software (Nedelec and Foethke, 2007). The simulation was performed in a 3D spherical geometry of 70 µm in diameter, corresponding to the average size of the starfish oocyte nucleus. Microtubules were modeled as dynamic, non-flexible polymers nucleated from the centrosomes. Centrosomes were static and positioned 3 µm from the cell cortex and 6 µm apart from each other (Fig. S6A). Microtubules touching the cell cortex underwent catastrophe immediately. No chromatin-mediated microtubule nucleation occurred in simulations and was neither observed experimentally (Fig. S6F). The inactivation of kinetochores to prevent early capture events was modeled by setting microtubule binding rate to 0 for the first 4 min after NEBD, and then increase it to 10-30-70-100%, every minute, 4-8 minutes after NEBD. To decrease computational costs, the simulation time-step was set to 0.05 s and microtubule rotation was neglected. Parameters of the model are tabulated on Fig. S6B. The configuration file to run simulations is available upon request. Cytosim is an Open Source project hosted on www.github/nedelec/cytosim.

## Acknowledgements

We are thankful to the members of the Lenart lab for reagents and support, in particular Michal Fleszar for assistance, and Joana Borrego-Pinto and Philippe Bun for sharing plasmid constructs and analysis tools. We are grateful to Serge Dmitrieff (Jacques-Monod Institute, Paris) for help with implementing Cytosim scripts. We thank Johanna Bischof (Tufts University, MA, USA), Natalia Wesolowska (EMBL) and Alexey Khodjakov (Wadsworth Center, NY, USA) for comments on the manuscript. We would like to additionally acknowledge the essential support of EMBL’s Advanced Light Microscopy Facility (ALMF), specifically Christian Tischer for the help with image analysis, IT support, and Laboratory Animal Resources (LAR), and Kresimir Crnokic (LAR), in particular. We also thank the EMBL Centre for Statistical Data Analysis and Bernd Klaus for the help with statistics, Stefano Maffini and Andrea Musacchio (MPI Dortmund, Germany) for mEGFP-Ndc80 protein, Beáta Bugyi (University of Pécs, Hungary) for Arp2/3-Alexa488 protein, and Rudolf Walczak and Iain Mattaj (EMBL) for the RanT24N protein.

The work was supported by the European Molecular Biology Laboratory, and specifically by the EMBL International PhD Programme and Darwin Trust of Edinburgh to M.B.

## Author contributions

M.B. and P.L. conceived the project and designed experiments. M.B. and A.C. performed experiments and analyzed data. M.B. and F.N. implemented and run simulations. M.B. and P.L. wrote the manuscript.

## Supplemental Material

### Supplemental Figures

**Supplemental figure S1. Dynein transports chromosomes along microtubules (A)** Maximum intensity z-projection from a time series of an oocyte expressing H2B-3mEGFP to label chromosomes. Ciliobrevin D was added 10 min before NEBD. Right: temporal color-coded maximum projection. Time is in mm:ss relative to NEBD. Scale bar: 10 µm. **(B)** Plot of chromosome distance to the AP over time in Ciliobrevin D treated oocytes. **(C)** Distances from the AP at NEBD for individual chromosomes from the analyzed set of control oocytes (shown on Fig. 2B). The plot reveals homogenous sampling of the data with regard to the distance to the AP. **(D)** Distances from the center of the nucleus at NEBD for individual chromosomes from the control dataset (shown on Fig 2B). The plot reveals homogeneous sampling with regard to the distance from the center of the nucleus.

**Supplemental figure S3. F-actin network does not interfere with chromosome capture and transport (A)** Single confocal slice from a time series of an oocyte expressing EB3-3mCherry to label microtubule plus-tips. Scale bar: 10 µm. Right: histogram showing the microtubule length distribution in DMSO control (red) and Latrunculin B treated (blue) oocytes. Thin lines: data from individual oocytes; thick lines: mean for 6 oocytes; shaded areas represent standard deviation. **(B)** Maximum intensity z-projection from a time series of an oocyte expressing H2B-3mEGFP to label chromosomes. Arrows indicate chromatin fragments resulting from the treatment with the DNA damaging agent, Zeocin. Scale bar: 10 µm. **(C)** Maximum intensity z-projection from a time series of an oocyte expressing H2B-3mEGFP to label chromosomes. Two minutes after NEBD the oocyte was injected with a pulse of Phalloidin-Alexa568 next to the nuclear area to stabilize filaments of the F-actin network. Scale bar: 10 µm. Time relative to NEBD in mm:ss. Right: plot of chromosome distance to the AP over time.

**Supplemental figure S5. Complete dataset for quantification of F-actin patch disassembly kinetics (A)** Selected maximum intensity z-projections from a 3D confocal time series though the oocyte’s nuclear region during chromosome congression for control and Latrunculin B treated oocytes shown on Fig. 5E-H. Chromosomes (H2B-3mEGFP) are in cyan and F-actin (3mCherry-UtrCH) in grey. Scale bars: 10 µm. **(B)** Left panels: plot of chromosome distance to the AP over time for control and Latrunculin B treated oocytes shown on **(A)**. Right panels: normalized 3mCherry-UtrCH intensity profile for chromosomes tracked on the left. Intensity is calculated in a 5 µm diameter sphere around the chromosome’s center of mass. Green: F-actin-driven, red: microtubule-driven transport, grey: chromosomes arrived to the AP. Chromosome capture events are represented as dots.

**Supplemental figure S6. Details of the computer simulations (A)** Renderings of the 3D computer simulation in Cytosim: the nucleus is represented as a sphere with a diameter of 70 µm. Chromosomes (cyan) with two kinetochores each (green) are transported by the contractile F-actin network. After capture of kinetochores by microtubules, chromosomes are transported by dynein to the centrosomes. **(B)** Table summarizing the main parameters used in the model. **(C)** Chromosome pair-wise velocity analysis to derive the network contraction rate, as described in (Bun et al., 2018). Comparison between experimental data (red) and simulation (grey). **(D)** Microtubule length distribution in oocytes (red) and simulations (black) fitted with the half-normal distribution function. Microtubule length was compared at time 3 to 10 min after NEBD, when asters reach full size. **(E)** Chromosome and kinetochore morphology in the experiment *versus* simulation. Left (experiment): spindle area during prometaphase, in an oocyte expressing H2B- 3mEGFP to label chromosomes and injected with mEGFP-Ndc80 protein to label kinetochores. Single deconvolved confocal slice. Scale bar: 2 µm. Right (simulation): chromosomes are represented as 1.6 µm spheres (blue) each with two 0.5 µm kinetochores located on the opposite sides. **(F)** Chromatinmediated microtubule nucleation is not active during the initial chromosome congression. Single deconvolved confocal slices through an oocyte fixed 8 minutes after NEBD and immunostained for tubulin (DM1α, grey), chromosomes (Draq5, cyan), and Phalloidin-A568 for F-actin (grey). Oocytes were treated prior maturation with Cytochalasin D, Nocodazole or DMSO as indicated.

## Supplemental Videos

**Video 1. After F-actin-driven transport, chromosomes form lateral attachments and are transported along the microtubules to the AP** Confocal sections taken every 0.7 s through the nuclear region of an oocyte expressing EB3- 3mCherry to visualize microtubule plus-tips (grey) and H2B-3mEGFP to label the chromosomes (cyan). Video starts at 06:25 after NEBD and runs for 8 minutes. Imaged area: 51.5×51.5 µm. Selected frames are shown in Fig. 1B.

**Video 2. F-actin and chromosome dynamics upon acute F-actin depolymerization** Maximum intensity z-projections through the nuclear region of live oocytes expressing H2B-3mEGFP to label chromosomes and 3mCherry-UtrCH to label F-actin. Latrunculin B, or corresponding amount of DMSO, was added at NEBD onset (00:00). Video starts at 00:30 after NEBD and runs for 23 minutes. Time step: 5 s. Imaged area: 94×94×70.5 µm. Still frames from this dataset are shown in Fig. 2B.

**Video 3. Chromosome capture is coordinated in F-actin dependent manner** Maximum intensity z-projections through the nuclear region of live oocytes expressing H2B-3mEGFP to label chromosomes. Latrunculin B, or corresponding amount of DMSO, was added at NEBD onset (00:00). Video starts at 00:30 after NEBD and runs for 17 minutes. Time step: 3 s. Imaged area: 68×68×60 µm. Still frames from this dataset are shown in Fig. 2C.

**Video 4. Chromosomes transported along microtubules are pulled through the F-actin network** Single confocal sections acquired every 1 s of the nuclear area of an oocyte expressing 3mEGFP-UtrCH (grey) and H2B-mCherry (cyan). Video starts at 15:00 after NEBD and runs for 3.4 min. Imaged area: 60×60 µm. Still frames from this dataset are shown in Fig. 3C.

**Video 5. F-actin patch disassembly kinetics correlates with chromosome capture by microtubules** Maximum intensity z-projections of z-stacks acquired every 13 s through the nuclear region of live oocyte expressing mEGFP-ArpC1 (grey), and H2B-mCherry (cyan). Video starts at NEBD onset (00:00) and runs for 15 min. Imaged area: 82×82×62 µm. Still frames from this dataset are shown in Fig. 5A.

**Video S6. Simulation of microtubule ‘search-and-capture’ in starfish oocytes** Video starts at 00:00 at NEBD and runs for 15 minutes. Still frames from this data set are shown in Fig. 6A (Model 3) and S6A. See the corresponding figure legends for details.

## References

Amiel, A., and E. Houliston. 2009. Three distinct RNA localization mechanisms contribute to oocyte polarity establishment in the cnidarian Clytia hemisphærica. Dev. Biol. 327:191–203. doi:10.1016/j.ydbio.2008.12.007.

Barisic, M., P. Aguiar, S. Geley, and H. Maiato. 2014. Kinetochore motors drive congression of peripheral polar chromosomes by overcoming random arm-ejection forces. Nat. Cell Biol. 16:1249–1256. doi:10.1038/ncb3060.

Borrego-Pinto, J., K. Somogyi, M.A. Karreman, J. König, T. Müller-Reichert, M. Bettencourt-Dias, P. Gönczy, Y. Schwab, and P. Lénárt. 2016a. Distinct mechanisms eliminate mother and daughter centrioles in meiosis of starfish oocytes. J. Cell Biol. 212:815–827. doi:10.1083/jcb.201510083.

Borrego-Pinto, J., K. Somogyi, and P. Lénárt. 2016b. Live Imaging of Centriole Dynamics by Fluorescently Tagged Proteins in Starfish Oocyte Meiosis. In Methods in molecular biology (Clifton, N.J.). 145–166.

Bun, P., S. Dmitrieff, J.M. Belmonte, F.J. Nédélec, and P. Lénárt. 2018. A disassembly-driven mechanism explains F-actin-mediated chromosome transport in starfish oocytes. Elife. 7. doi:10.7554/eLife.31469.

Burkel, B.M., G. Von Dassow, and W.M. Bement. 2007. Versatile fluorescent probes for actin filaments based on the actin-binding domain of utrophin. Cell Motil. Cytoskeleton. 64:822–832. doi:10.1002/cm.20226.

Cai, S., C.B. O’Connell, A. Khodjakov, and C.E. Walczak. 2009. Chromosome congression in the absence of kinetochore fibres. Nat. Cell Biol. 11:832–838. doi:10.1038/ncb1890.

Carazo-salas, R.E., and E. Karsenti. 2003. Long-Range Communication between Chromatin and Microtubules in Xenopus Egg Extracts. Curr. Biol. 13:1728–1733. doi:10.1016/j.

Chesarone, M.A., A.G. DuPage, and B.L. Goode. 2010. Unleashing formins to remodel the actin and microtubule cytoskeletons. Nat. Rev. Mol. Cell Biol. 11:62–74. doi:10.1038/nrm2816.

Crowder, M.E., M. Strzelecka, J.D. Wilbur, M.C. Good, G. von Dassow, and R. Heald. 2015. A Comparative Analysis of Spindle Morphometrics across Metazoans. Curr. Biol. 25:1542–1550. doi:10.1016/j.cub.2015.04.036.

Dasso, M., T. Seki, Y. Azuma, T. Ohba, and T. Nishimoto. 1994. A mutant form of the Ran/TC4 protein disrupts nuclear function in Xenopus laevis egg extracts by inhibiting the RCC1 protein, a regulator of chromosome condensation. EMBO J. 13:5732–44.

Dehapiot, B., V. Carrière, J. Carroll, and G. Halet. 2013. Polarized Cdc42 activation promotes polar body protrusion and asymmetric division in mouse oocytes. Dev. Biol. 377:202–212. doi:10.1016/j.ydbio.2013.01.029.

Deng, M., P. Suraneni, R.M. Schultz, and R. Li. 2007. The Ran GTPase Mediates Chromatin Signaling to Control Cortical Polarity during Polar Body Extrusion in Mouse Oocytes. Dev. Cell. 12:301–308. doi:10.1016/j.devcel.2006.11.008.

Hayden, J.H., S.S. Bowser, and C.L. Rieder. 1990. Kinetochores capture astral microtubules during chromosome attachment to the mitotic spindle: Direct visualization in live newt lung cells. J. Cell Biol. 111:1039–1045. doi:10.1083/jcb.111.3.1039.

Heald, R., and A. Khodjakov. 2015. Thirty years of search and capture: The complex simplicity of mitotic spindle assembly. J. Cell Biol. 211:1103–1111. doi:10.1083/jcb.201510015.

Heald, R., R. Tournebize, T. Blank, R. Sandaltzopoulos, P. Becker, A. Hyman, and E. Karsenti. 1996. Self-organization of microtubules into bipolar spindles around artificial chromosomes in Xenopus egg extracts. Nature. 382:420–425. doi:10.1038/382420a0.

Henty-Ridilla, J.L., A. Rankova, J.A. Eskin, K. Kenny, and B.L. Goode. 2016. Accelerated actin filament polymerization from microtubule plus ends. Science (80-.). 352:1004–1009. doi:10.1126/science.aaf1709.

Hetzer, M., O.J. Gruss, and I.W. Mattaj. 2002. The Ran GTPase as a marker of chromosome position in spindle formation and nuclear envelope assembly. Nat. Cell Biol. 4:E177–E184. doi:10.1038/ncb0702-e177.

Holy, T.E., and S. Leibler. 1994. Dynamic instability of microtubules as an efficient way to search in space. Proc. Natl. Acad. Sci. U. S. A. 91:5682–5.

Jaffe, L.A., and M. Terasaki. 2004. Quantitative microinjection of oocytes, eggs, and embryos. Methods Cell Biol. 74:219–42.

Kaksonen, M., Y. Sun, and D.G. Drubin. 2003. A pathway for association of receptors, adaptors, and actin during endocytic internalization. Cell. 115:475–87.

Kaláb, P., A. Pralle, E.Y. Isacoff, R. Heald, and K. Weis. 2006. Analysis of a RanGTP-regulated gradient in mitotic somatic cells. Nature. 440:697–701. doi:10.1038/nature04589.

Kalinina, I., A. Nandi, P. Delivani, M.R. Chacón, A.H. Klemm, D. Ramunno-Johnson, A. Krull, B. Lindner, N. Pavin, and I.M. Tolić-Nørrelykke. 2013. Pivoting of microtubules around the spindle pole accelerates kinetochore capture. Nat. Cell Biol. 15:82–87. doi:10.1038/ncb2640.

Kapoor, T.M., M.A. Lampson, P. Hergert, L. Cameron, D. Cimini, E.D. Salmon, B.F. McEwen, and A. Khodjakov. 2006. Chromosomes can congress to the metaphase plate before biorientation. Science (80-.). 311:388–391. doi:10.1126/science.1122142.

Karsenti Newport,J., Huble, R., E., and M. Kirschner. 1984. Interconversion of metaphase and interphasemicrotubule arrays, as studied by the injection of centrosomesand nuclei i nto Xenopus eggs. J.Cell.Biol. 98:1730–1745.

Kirschner, M.W., and T.J. Mitchison. 1986. Beyond self assembly: from microtubules to morphogenesis. Cell. 45:329–342.

Kolano, A., S. Brunet, A.D. Silk, D.W. Cleveland, and M.-H. Verlhac. 2012. Error-prone mammalian female meiosis from silencing the spindle assembly checkpoint without normal interkinetochore tension. Proc. Natl. Acad. Sci. U. S. A. 109:E1858-67. doi:10.1073/pnas.1204686109.

Lénárt, P., C.P. Bacher, N. Daigle, A.R. Hand, R. Eils, M. Terasaki, and J. Ellenberg. 2005. A contractile nuclear actin network drives chromosome congression in oocytes. Nature. 436:812–818.

Lénárt, P., and J. Ellenberg. 2002. Nuclear envelope dynamics in oocytes: From germinal vesicle breakdown to mitosis. Curr. Opin. Cell Biol. 15:88–95.

Li, Y., W. Yu, Y. Liang, and X. Zhu. 2007. Kinetochore dynein generates a poleward pulling force to facilitate congression and full chromosome alignment. Cell Res. 17:701–712. doi:10.1038/cr.2007.65.

Lowery, L.A., and D. Van Vactor. 2009. The trip of the tip: understanding the growth cone machinery. Nat. Rev. Mol. Cell Biol. 10:332–43. doi:10.1038/nrm2679.

Magidson, V., C.B. O’Connell, J. Lončarek, R. Paul, A. Mogilner, and A. Khodjakov. 2011. The Spatial Arrangement of Chromosomes during Prometaphase Facilitates Spindle Assembly. Cell. 146:555–567. doi:10.1016/j.cell.2011.07.012.

Magidson, V., R. Paul, N. Yang, J.G. Ault, C.B. O’Connell, I. Tikhonenko, B.F. Mcewen, A. Mogilner, and A. Khodjakov. 2015. Adaptive changes in the kinetochore architecture facilitate proper spindle assembly. Nat. Cell Biol. 17:1134–1144. doi:10.1038/ncb3223.

Mitchison, T., and M. Kirschner. 1984. Dynamic instability of microtubule growth. Nature. 312:237–242. doi:10.1038/312237a0.

Mogessie, B., and M. Schuh. 2017. Actin protects mammalian eggs against chromosome segregation errors. Science (80-.). 357:eaal1647. doi:10.1126/science.aal1647.

Monnier, N., S.-M. Guo, M. Mori, J. He, P. Lénárt, and M. Bathe. 2012. Bayesian Approach to MSD-Based Analysis of Particle Motion in Live Cells. Biophys. J. 103:616–626. doi:10.1016/j.bpj.2012.06.029.

Mori, M., N. Monnier, N. Daigle, M. Bathe, J. Ellenberg, and P. Lénárt. 2011. Intracellular transport by an anchored homogeneously contracting F-actin meshwork. Curr. Biol. 21:606–611. doi:10.1016/j.cub.2011.03.002.

Mori, M., K. Somogyi, H. Kondo, N. Monnier, H.J. Falk, P. MacHado, M. Bathe, F. Nédélec, and P. Lénárt. 2014. An Arp2/3 nucleated F-actin shell fragments nuclear membranes at nuclear envelope breakdown in starfish oocytes. Curr. Biol. 24:1421–1428. doi:10.1016/j.cub.2014.05.019.

Moss, D.K., A. Wilde, and J.D. Lane. 2009. Dynamic release of nuclear RanGTP triggers TPX2- dependent microtubule assembly during the apoptotic execution phase. J. Cell Sci. 122:644–655. doi:10.1242/jcs.037259.

Nedelec, F., and D. Foethke. 2007. Collective Langevin dynamics of flexible cytoskeletal fibers. New J. Phys. 9:427–427. doi:10.1088/1367-2630/9/11/427.

Petry, S., and R.D. Vale. 2015. Microtubule nucleation at the centrosome and beyond. Nat. Cell Biol. 17:1089–1093. doi:10.1038/ncb3220.

Prodon, F., J. Chenevert, and C. Sardet. 2006. Establishment of animal-vegetal polarity during maturation in ascidian oocytes. Dev. Biol. 290:297–311. doi:10.1016/j.ydbio.2005.11.025.

Rieder, C.L., and S.P. Alexander. 1990. Kinetochores are transported poleward along a single astral microtubule during chromosome attachment to the spindle in newt lung cells. J. Cell Biol. 110:81–95. doi:10.1083/jcb.110.1.81.

Schindelin, J., I. Arganda-Carreras, E. Frise, V. Kaynig, M. Longair, T. Pietzsch, S. Preibisch, C. Rueden, S. Saalfeld, B. Schmid, J.-Y. Tinevez, D.J. White, V. Hartenstein, K. Eliceiri, P. Tomancak, and A. Cardona. 2012. Fiji: an open-source platform for biological-image analysis. Nat. Methods. 9:676–682. doi:10.1038/nmeth.2019.

Shao, H., R. Li, C. Ma, E. Chen, and X.J. Liu. 2013. Xenopus oocyte meiosis lacks spindle assembly checkpoint control. J. Cell Biol. 201:191–200. doi:10.1083/jcb.201211041.

Shrestha, R.L., and V.M. Draviam. 2013. Lateral to End-on Conversion of Chromosome-Microtubule Attachment Requires Kinesins CENP-E and MCAK. Curr. Biol. 23:1514–1526. doi:10.1016/j.cub.2013.06.040.

Webster, A., and M. Schuh. 2017. Mechanisms of Aneuploidy in Human Eggs. Trends Cell Biol. 27:55–68. doi:10.1016/j.tcb.2016.09.002.

Wollman, R., E.N. Cytrynbaum, J.T. Jones, T. Meyer, J.M. Scholey, and A. Mogilner. 2005. Efficient chromosome capture requires a bias in the “search-and-capture” process during mitotic-spindle assembly. Curr. Biol. 15:828–832. doi:10.1016/j.cub.2005.03.019.

Yamagishi, Y., and H. Abe. 2017. Actin assembly mediated by a nucleation promoting factor WASH is involved in MTOC-TMA formation during Xenopus oocyte maturation. Cytoskeleton. doi:10.1002/cm.21428.

Yang, Z., U.S. Tulu, P. Wadsworth, and C.L. Rieder. 2007. Kinetochore Dynein Is Required for Chromosome Motion and Congression Independent of the Spindle Checkpoint. Curr. Biol. 17:973–980. doi:10.1016/j.cub.2007.04.056.

